# Recipient and donor characteristics govern the hierarchical structure of heterospecific pollen competition networks

**DOI:** 10.1101/2020.01.21.914515

**Authors:** Jose B. Lanuza, Ignasi Bartomeus, Tia Lynn Ashman, Romina Rader

## Abstract

- Pollinator sharing can have negative consequences for plant fitness with the arrival of foreign pollen, yet responses are often variable among species. Plant traits and relatedness of donor and recipient species have been suggested to drive the variations in plant fitness, but how they shape the structure of pollen competition networks has been overlooked at the community level.
- To understand the importance of traits and relatedness we conducted a controlled glasshouse experiment with an artificial co-flowering community. We performed 900 reciprocal crosses by experimentally transferring pollen among 10 species belonging to three different plant families.
- We found a significant reduction in seed set for 67% of the crosses, driven largely by recipient traits and the interaction between recipient-donor traits under specific circumstances of trait-matching. These traits and their asymmetries among species led to a hierarchical (or transitive) structure of pollen competition with clear winners and losers depending on specific combination of traits.
- A greater understanding of the importance of trait matching and asymmetries among donor and recipient plant species will facilitate knowledge of the mechanisms underlying foreign pollen impacts upon plant reproductive fitness. This will require a shift from pairwise to community level interactions.

## INTRODUCTION

In natural ecosystems, several plant species coexist simultaneously and share floral visitors (Waser *et al.*, 1996; Carvalheiro et al., 2014). From the plants’ perspective, pollinator sharing can be positive if it increases overall pollinator populations, as an increase in the number of visits often correlates with a greater probability of fertilization (Engel & Irwin, 2003). Yet, among these possible flower visitors there are also inefficient pollinators (Inouye, 1980; Magrach *et al.*, 2017) and inconstant pollinators that transfer foreign pollen from other plant species. In fact, many pollinators are responsible for conspecific pollen loss and the transport of foreign pollen to plants, both of which can have important detrimental effects on plant fitness (Morales & Traveset, 2009; Ashman & Arceo-Gómez, 2013; Moreira-Hernández & Muchhala, 2019).

Foreign pollen arrival can play an important role in plant fitness but outcomes are variable and appear to be context dependent (Arceo-Gómez *et al.*, 2019). Some of this variation is likely due to the enormous variability on the amount of foreign pollen transferred across systems, ranging from 0 to 75 percent of total pollen deposited on floral stigmas (Fang *et al.*, 2019). While most studies report average heterospecific pollen loads of less than 20 percent (Bartomeus *et al.*, 2008; Ashman & Arceo-Gómez, 2013; Fang & Huang, 2013), even low amounts of heterospecific pollen transferred can significantly decrease fitness (Thomson *et al.*, 1982; Loughnan *et al.*, 2014).

Although we have some understanding of the impacts of heterospecific pollen quantity, where greater heterospecific pollen loads are expected to cause a greater impact (Harder *et al.*, 1993; Briggs et al., 2015), we have little knowledge of the relevance of donor and recipient identity upon plant fitness (Arceo-Gómez *et al.*, 2019). Floral traits are thought to be important in mediating the impact of foreign pollen transfer on plant fitness (Ashman & Arceo-Gómez, 2013). However, the multifactorial nature of functional traits and their interactions among donor and recipient species across multiple plant families in a co-flowering community make it difficult to determine exactly which traits or combination of traits are potentially driving the observed effects.

Pollen size, pollen quantity, number of apertures and allelopathy are thought to be key components in understanding the outcome of foreign pollen arrival (Murphy & Aarssen, 1995b; Ashman & Arceo-Gómez, 2013). For example, small pollen is predicted to decrease seed production because it is more likely to clog the opening of the stigmatic surface. Yet, large pollen can outcompete smaller pollen grains due to faster pollen tube growth rates (Williams & Rouse, 1990). In addition, heterospecific pollen arrival is positively correlated with stigma size (Montgomery & Rathcke, 2012), and therefore species with larger stigmas could be more vulnerable to heterospecific pollen. The trait-matching between donor and recipient species is also likely to impact plant fitness (Arceo-Gómez *et al.*, 2019). For instance, larger pollen grains could potentially clog smaller stigmas with fewer pollen grains, and broader stigmas are less likely to be clogged by smaller pollen grains (Galen & Gregory, 1989). Further, species mating systems are also thought to play a key role in the species response to heterospecific pollen, where self-incompatible species with self-pollen recognition could have the ability to exclude foreign pollen (McClure *et al.*, 2011). Moreover, the competitive ability of pollen of these self-incompatible species is also likely to be greater due to their outcrossing nature (Ashman & Arceo-Gómez, 2013). Yet, we still do not have a clear picture of how these traits and trait-matching combinations determine the impact of heterospecific pollen receipt.

Closely related species are more likely to have similar traits (Letten & Cornwell, 2015). Similar traits can result in a higher probability that heterospecific pollen will interact with ovules, and hence greater negative effects are thought to be associated with more closely related species, i.e., congeners (Arceo-Gómez & Ashman, 2011 Brown & Mitchell, 2001; Tong & Huang, 2016). However, less attention has been given to the effect of heterospecific pollen on distantly related species- i.e., different order or family (Galen & Gregory, 1989; Neiland & Wilcock, 1999). Given that pollen carried on many insects and stigmas belongs to multiple species with differing degrees of relatedness (Arceo-Gómez & Ashman, 2016; Fang & Huang, 2013), understanding the effect of heterospecific pollen of close and distantly related species requires a systematic approach to disentangle the importance of relatedness at community level.

Identifying the mechanisms driving foreign pollen impacts is challenging as experimental studies to date have largely focused on species pairs. Yet, evidence is mounting that variations in trait matching and phylogeny are important in mediating heterospecific impacts upon plant fitness. For instance, Tong & Huang (2016) demonstrate an asymmetrical effect of heterospecific pollen in 6 species of *Pedicularis* whereby foreign pollen of long-styled species was able to grow the full length of the style on short-styled species, but not vice-versa. If such asymmetrical effects exist, there are likely winners and losers within co-flowering communities that govern the structure of the effects of heterospecific pollen competition networks. However, the structure of how different plant species compete at the community level remains unexplored.

Pollinator interactions between donor and recipient co-flowering plants will govern the competitive outcome of heterospecific pollen deposition. The combination of species interacting with each other could result in a structure that is hierarchical (i.e. with clear winners and losers) or non-hierarchical where there are no clear winners or losers, akin to the rock-paper-scissor game (Kerr *et al.*, 2002). Intransitive or non-hierarchical structures are thought to be a possible mechanism of species coexistence where there is not a clear dominant species (Soliveres & Allan, 2018) whereby a hierarchical or transitive structure can be destabilizing when the most dominant species exclude the rest (Alcántara *et al.*, 2017). Despite the increasing interest in interspecific pollen competition, the structure of donor and recipient networks in co-flowering communities remains unknown, yet is critical to inform the potential for plant interspecific pollen to contribute to species coexistence at the community scale.

Here, we investigate how male and female plant reproductive traits and relatedness of donor and recipient species mediate the impact of heterospecific pollen and the structure of the interspecific pollen competition network by creating an artificial co-flowering community in a glasshouse with 10 species belonging to three different families that varied in reproductive traits. Our study addressed the following questions:

1. To what extent does heterospecific pollen receipt impact plant reproductive fitness?
2. Is the structure of the pollen competition network hierarchical or non-hierarchical?
3. How do phylogenetic distance and plant-pistil or pollen traits mediate the impacts of heterospecific pollen receipt on seed set?

## MATERIALS AND METHODS

### Experimental design

The study was conducted at University of New England (Armidale, Australia) from November 2017 to March 2018. The experimental design comprised 10 plant species (N=20 for each species) from three different families: Brassicaceae (4), Convolvulaceae (2) and Solanaceae (4) (Supporting Information, **Table S1**). The species of the study had contrasting reproductive traits, including pistil and pollen traits, and different mating systems (see **Table S2** and **Fig. S1** for traits and their correlations). Moreover, the selected species had also different degree of relatedness (see phylogenetic tree, **Fig. 1**). Hence, the reciprocal crosses between all species allowed us to test multiple different combinations between species with a wide range of traits and relatedness. The criteria for species selection were: 1) variation in traits and relatedness, 2) low structural flower complexity for ease of pollination and 3) fast life cycle and simultaneous flowering. All the species served as pollen recipients and donors. Species were watered once or twice per day and fertilized weekly (NPK 23: 3.95: 14) and the rooms of the glasshouse were temperature controlled with temperature oscillations between day and night (28 to 22 °C).

**Fig. 1.**
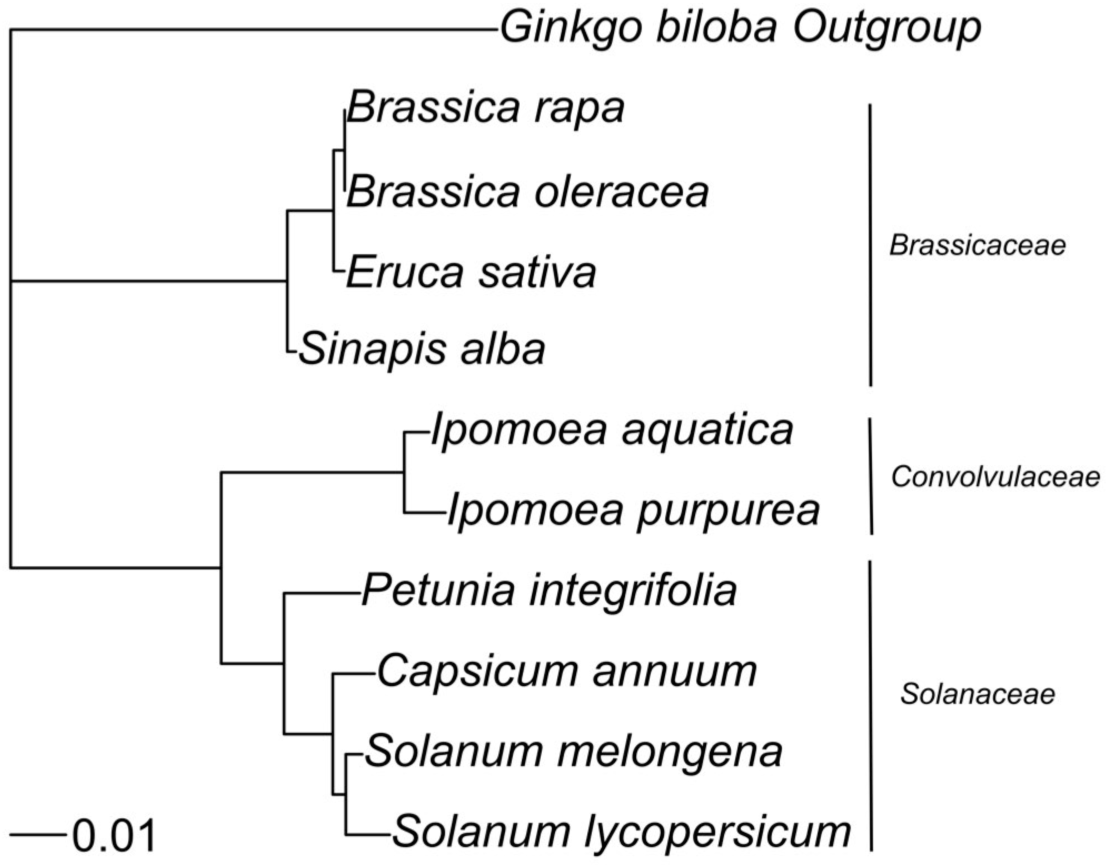
Phylogenetic tree of the ten different species with a systematic design to have a wide range of relatedness for the different reciprocal heterospecific pollen treatments. From top to bottom the families are Brassicaceae, Convolvulaceae and Solanaceae with *Ginkgo biloba* as an outgroup.

### Species reproductive biology

Different mating strategies can lead to tolerance or vulnerability of heterospecific pollen (Ashman & Arceo-Gómez, 2013), for this reason we tested the reproductive biology of the species with the following pollination treatments: 1) hand cross pollination (between individuals of the same species), 2) hand self pollination, 3) apomixis (bagged emasculated flowers) and 4) natural selfing (untouched bagged flowers). We performed 10 replicates per treatment and recorded seed set as the proportion of ovules converted to seeds for all of them.

### The impact of heterospecific pollen receipt on seed set

The impact of foreign pollen was evaluated using two different types of heterospecific pollen loads (i) 50% conspecific pollen and 50% heterospecific pollen and (ii) 100% foreign pollen to determine if foreign pollen can trigger fruit/seed production or hybridization in closely related species. For each recipient species, we performed 18 different treatments, 9 treatments with 50% heterospecific pollen and 9 treatments with 100% heterospecific pollen. Therefore, we performed 180 treatments with 10 replicates per treatment resulting in a total of 1800 pollination events. Seed production per flower was the effect indicator for all treatments. For the treatments with foreign pollen and hand cross pollination, flowers were emasculated the day prior to anthesis and hand pollinated the next day. Hand pollination was conducted with 3-4 brushes of a toothpick from an Eppendorf tube onto the stigma surface. For each species, 20 anthers were collected and their pollen counted with a hemocytometer. We counted the pollen within each anther 4 times, averaged these counts and calculated the average of 20 anthers. Anthers were squashed on 1 ml of distilled water and homogenized in a vortex for 30 seconds. Using the average number of pollen grains per anther we calculated the proportion of anthers per mix to achieve a 50-50% mix. The anthers were mixed in Eppendorf tubes of 1.5 ml and vibrated with an electric toothbrush (with head of toothbrush removed) under the outside of the tube. For the species requiring buzz-pollination, *S. lycopersicum* and *S. melongena*, we used the same procedure with an electric toothbrush at the bottom of the corolla/receptacle to extract the pollen directly into the tubes. For the rest of the species the anthers were placed directly into the tubes. To confirm that the treatments achieved the desired ratio, the total stigmatic load of pollen was counted from additional stigmas collected after 10 hours of the hand pollination treatment (**Fig. S2**). We squashed the stigmas onto a slide with a grid and all of the pollen on the stigma was counted using a microscope (40x magnification). Moreover, we also calculated the proportions of pollen in this 50% pollen mix per species and family (**Fig. S3; Fig. S4**). We counted two replicate 50% pollen treatments for each donor species from a different family from the focal species (20 counts in total). For example, for the species *S. alba* (Brassicaceae) we counted one treatment (50% heterospecific pollen) with one Solanaceae (*S. melongena*) and one Convolvulaceae (*I. aquatica*). We did not count heterospecific pollen mixes between species of the same families because we expected them to have similar properties (e.g., size, stickiness).

We tested the accuracy of our treatments by comparing pollen proportions among families with linear models and a posthoc Tukey test (Supporting Information **Fig. S5**), function *lm* of the R stats package version 3.5.1 and *emmeans* package version 1.3.5.1 (Lenth & Lenth, 2018). Moreover, we tested the correlation between the total amount of pollen deposited on the stigma and stigma size in the heterospecific treatments through Pearson’s correlation with the R base function from the R stats package *cor.test* to determine whether differences in effect could be explained by pollen quantity (Pearson’s correlation = 0.57, *P* < 0.01). Finally, we used as supporting analysis of our methodology a linear model with effect sizes as response variable and pollen ratios as fixed effect (*HP effect size* ∼ *Pollen ratios*), because we have just a subset of 20 combinations of recipient x donor pollen counts, we tested exclusively for this subset and we did not include pollen ratios in the main model of absolute effect of traits. We found non-significant changes in effect sizes due to pollen ratios as fixed effects (Sum of squares = 0.46, *F*-value = 0.39, *P* = 0.54).

To evaluate the overall effect of 50% heterospecific pollen treatments on seed production we performed linear models (*lm* function from the R stats package version 3.5.1) of the log transformed seed set data. For each species the seed production with the 50% heterospecific pollen treatments was compared with the average seed production of hand cross pollination (just conspecific pollen) by releveling it as a reference level.

### Heterospecific pollen network competition structure

To compare the effect of heterospecific pollen across species we calculated standardized Hedges’ g [(mean seed set 50% Heterospecific Conspecific mix - mean seed set 100% Conspecific)/pooled SD] with the *effsize* package version 0.7.4 (Torchiano, 2016). We calculated Hedges’ g at three different levels to determine the impact of heterospecific pollen on seed production: (i) we determined the effect sizes of each treatment per species (i.e. recipient species x donor species matrix) which gives information of the individual donors on each focal species; (ii) we also calculated the effect sizes for each focal species at donor family level (i.e. recipient species x donor family matrix) to see if there are differences in effect within the three families; and (iii) we estimated the grouped effect of all donors per focal species (i.e. recipient species x grouped effect donor matrix) which emphasizes overall recipient species response to heterospecific pollen.

We analyzed the network structure of heterospecific pollen by converting the species-species matrix of effect sizes into a dominance matrix (binary matrix). For this, we considered that species A is a “winner” (1) and B a “loser” (0) when the effect size of A as a recipient is smaller than the effect size of B as a recipient. This means that A+B on A are causing comparatively lower reduction of seed set than A+B on B. We determined the structure or hierarchy of heterospecific pollen competition through the triangle transitivity (*T _tri_*) method described in Shizuka & McDonald (2012). This approach has been already used in different fields such as network theory (Lerner & Lomi, 2017), animal behavior (Nandi *et al.*, 2014; de Silva, Schmid, & Wittemyer, 2017; de Silva *et al.*, 2017) and species competition (Godoy *et al.*, 2017) among others. Following this method, the dominance matrix is converted to a network format with the help of *statnet* package version 2018.10 (Handcock *et al.*, 2008) and then divided in all the possible combinations between three species (triads). We calculated the relative proportion of transitive triads with the help of *sna* package version 2.4 (Butts, 2008). There are two possible type of triads, transitive or hierarchical where one species dominates over the rest A>B>C and intransitive or non-hierarchical where there is not a clear winner in the triad A > B, B > C, C > A (Appleby, 1983). We calculated the relative proportion of transitive triads for a total of 120 triplets and then obtained a *P*-value through randomization test by comparing with 1000 simulations with the same number of triads (Shizuka & McDonald 2012). Finally, we represented graphically the dominance matrix converted to network by using the package *igraph* (1.2.4) (Csárdi & Nepusz, 2006).

### Phylogenetic distance

We conducted Procrustes analysis to check for correlations between the matrix of effect sizes (species x species matrix) and the matrix of phylogenetic distance. To calculate the phylogenetic distances among the species we downloaded the sequences of the Internal transcribed spacer (ITS) and ribulose-bisphosphate carboxylase (RBCL) gene from GenBank (https://www.ncbi.nlm.nih.gov/genbank/, accessed 20 Oct. 2018). The sequences were then aligned with the online software Clustal Omega and the pairwise evolutive distances calculated with MEGA7 for both markers. We calculated the mean phylogenetic distance from both markers. In addition, we used the square root of the evolutive distance as this is thought to be a stronger approach to understand the displacement between two taxa rather than assuming simple linear divergence (Letten & Cornwell 2015). Finally, we tested the phylogenic signal of traits with all the species. We calculated the phylogenetic signal with the package phytools version 0.6-60 (Revell, 2012) where Pagel’s lambda was used as a measurement of phylogenetic signal.

### Plant functional traits

The traits measured for each species were: 1) pollen per anther, 2) pollen size, 3) number of ovules, 4) pollen-ovule ratio, 5) ovary length, 6) ovary width, 7) style length, 8) style width, 9) stigma length, 10) stigma width, 11) stigmatic area, 12) selfing rate and 13) self-incompatibility index (see **Table S2** for traits and their values). All the morphometrical measurements were performed with a stereo-photomicroscope with the exception of pollen size that was carried out with a light microscope. Ovule number was counted with the help of a stereomicroscope and a small grid over a petri dish from 15 randomly selected flowers. For all the species we counted the number of seeds produced per flower treated. Selfing rate was calculated as the percentage of ovules that were converted to seeds after self pollination. Levels of self-incompatibility were estimated for each species by dividing the seed set of hand self pollination by hand cross pollination (Lloyd & Schoen, 1992).

First, in order to test the relative effect of traits (i.e. trait similarity) on seed production with foreign pollen we performed Procrustes analysis with symmetric rotation (*vegan* Package version 2.5-4; Oksanen *et al.*, 2019) between the species x species matrix of effect sizes and the distance matrix of all traits (Euclidean distances). Furthermore, we tested for correlations between the matrix of effect sizes and the different distance matrices of each trait separately (Euclidean distances). Because trends or correlations at family level can be obscured when considered across all the three families, we also undertook the same analysis for each family independently (separated traits and grouped traits) with the exception of the family Convolvulaceae which comprised too few species (n = 2).

Second, we analyzed the absolute effect of traits on the heterospecific pollen effect (effect sizes) through linear models (*lm* function from the R stats package version 3.5.1), where we tested the effect of different donor and recipient traits and their interaction on the seed set of the different recipient species. We created a unique model based on two different criteria, first we filtered from the available trait variables those that can be relevant in the understanding of heterospecific pollen effect based on our knowledge and previous literature, and second, due to high correlation among fixed effects, we simplified the model until we were able to have non-correlated fixed variables while maintaining the logic of the ecological questions. Therefore, we created the following model: *HP effect size* ∼ *Donor pollen size* * *Recipient stigma siz*e + *Recipient pollen-ovule ratio*. Finally, in order to compare the importance of recipient and donor identity on effect sizes we performed an additional linear model where recipient and donor identity were considered as single fixed effect (*HP effect size* ∼ *Donor identity* + *Recipient identity*) and then their importance on effect sizes was compared with the help of the function *anova* from the R stats package version 3.5.1.

## RESULTS

### Species reproductive biology

Species within Solanaceae and Convolvulaceae were highly self-compatible species with the exception of *P. integrifolia* (Solanaceae) which was only partially self-compatible (**Fig. S6**, **Table S3**). Species within Brassicaceae were highly self-incompatible in general, but there was some evidence of self-compatibility in *E. sativa*, and fully self-compatibility in *S. alba*. None of the remaining species produced seeds via apomixis or agamospermy and we found high levels of autonomous selfing for species within both Convolvulaceae and the Solanaceae *S. lycopersicum* and *S. melongena*.

### The impact of heterospecific pollen receipt on seed set

Heterospecific pollen treatments significantly reduced seed set for 67% of the 900 pairwise crosses in comparison with the control (i.e. hand cross pollination; **Tables S4-S5**). Seed production was negligible in the 100% heterospecific pollen treatments (**Table S6**).

The effect sizes of the different donors on each focal species (species x species matrix) and each family (species x family donor matrix) were quite homogeneous (**Fig 2; Fig. S7**). When we compared the grouped effect sizes of all the donors by focal species (**Fig. 3**), we see that all Solanaceae species in addition to *I. aquatica*, *B. oleracea* and *B. rapa* were highly susceptible to heterospecific pollen with large effect sizes. In contrast, *I. purpurea*, *E. sativa* and *S. alba* had a small to null effect sizes and therefore little impact of heterospecific pollen on their fitness. We found that the species had very different responses even within families. For instance, in the Solanaceae family, *C. annuum* and *S. lycopersicum* had twice the effect size of their confamilials *P. integrifolia* and *S. melongena.* The same pattern was seen for the remaining two families where *I. aquatica* (Convolvulaceae), *B. oleracea* (Brassicaceae) and *B. rapa* (Brassicaceae) had a large effect size and in contrast *I. purpurea* (Convolvulaceae), *S. alba* (Brassicaceae) and *E. sativa* (Brassicaceae) had small to a null effect.

**Fig. 2.**
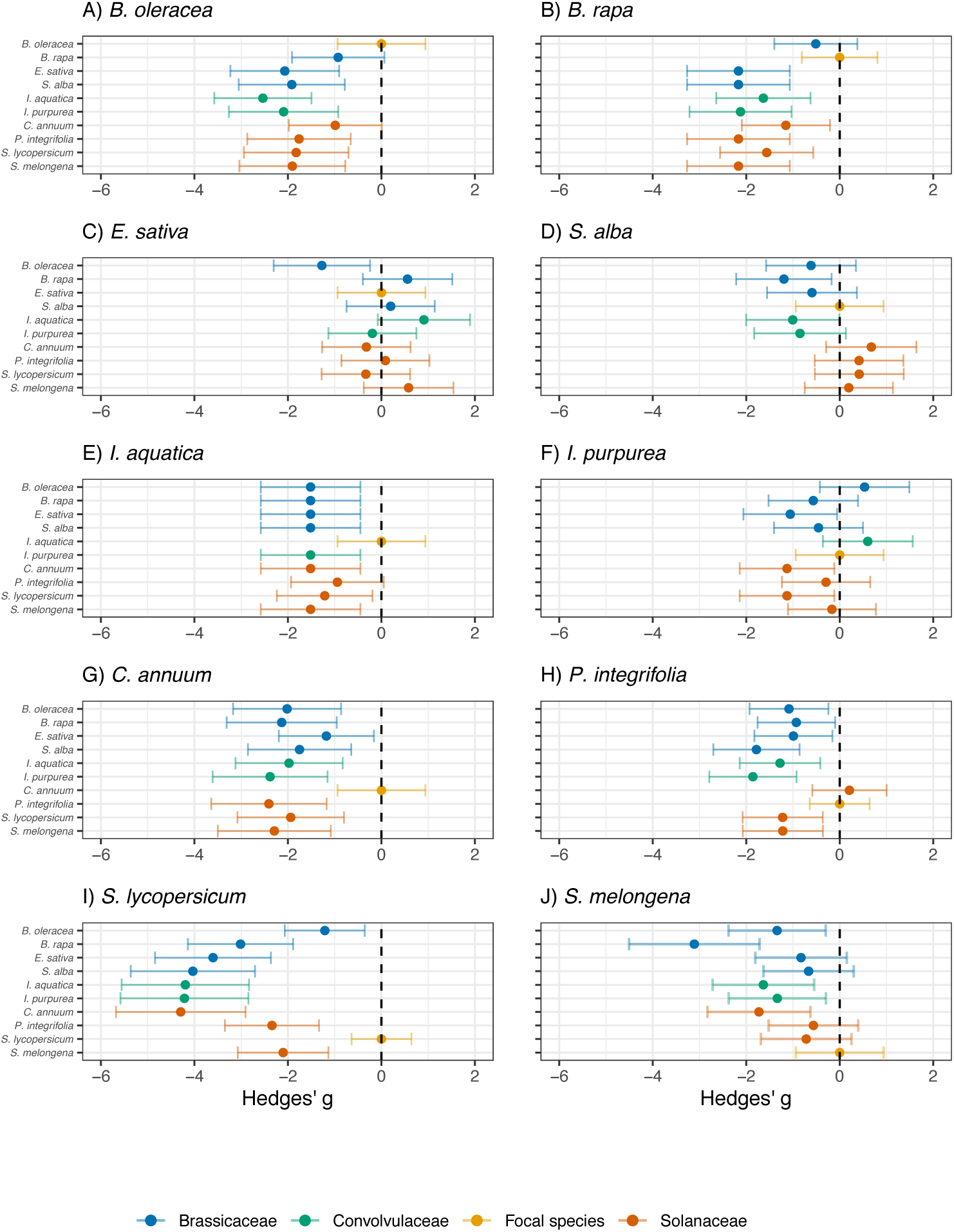
Effect sizes (95% confidence intervals) of the different nine treatments (50-50% pollen) on each focal species (recipient) and a hand cross pollination treatment (control). For each plot, the distinct families are coloured differently, Brassicaceae (blue), Convolvulaceae (green) and Solanaceae (orange), and the control, hand cross pollination with conspecific pollen (yellow). The dashed vertical line on 0 represents no effect.

**Fig. 3.**
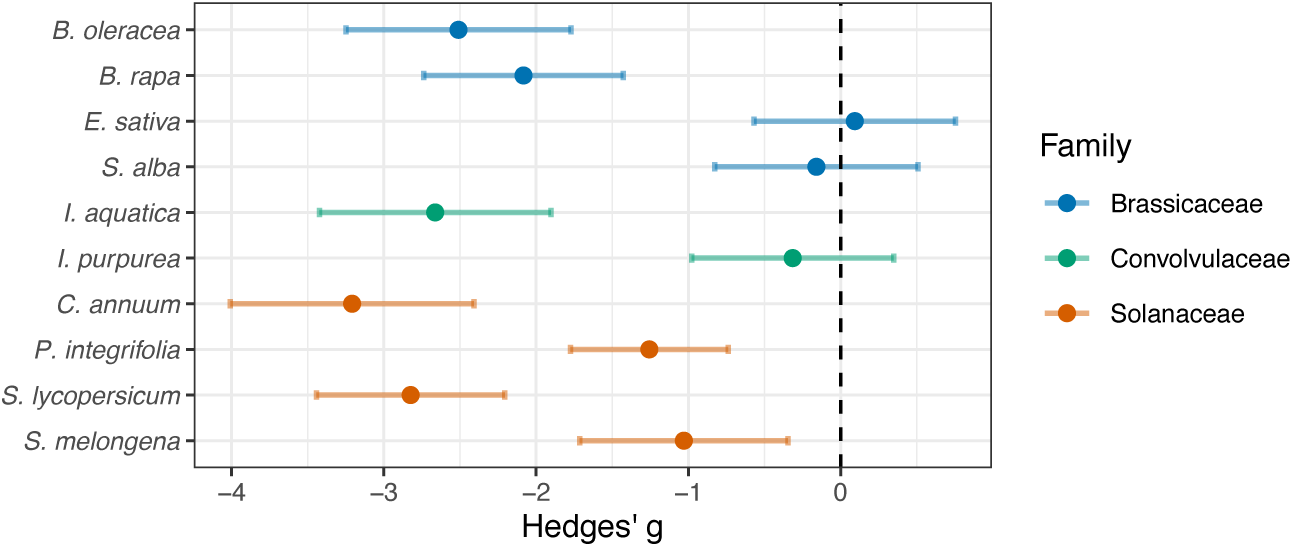
The impact of foreign pollen on recipient plant species. Grouped effect sizes (with 95% confidence intervals) of 9 different donor species of heterospecific pollen upon all recipients. The effect sizes of the different species are coloured by family, Brassicaceae (blue), Convolvulaceae (green) and Solanaceae (orange). The dashed vertical line on 0 represents no effect.

### Heterospecific pollen network competition structure

We found that out of the 120 possible triads (combinations of three species), 117 (97.5%) triads were transitive and just 3 (2.5%) intransitive. The proportion of intransitive triangles or triangles with cyclic competition (where there is not a clear winner or dominant species) was less than expected when we compared with the random simulated networks (*P* < 0.001). Therefore, the structure of pollen competition under experimental conditions of pollen transfer is highly transitive or hierarchical (**Fig. 4**).

**Fig. 4.**
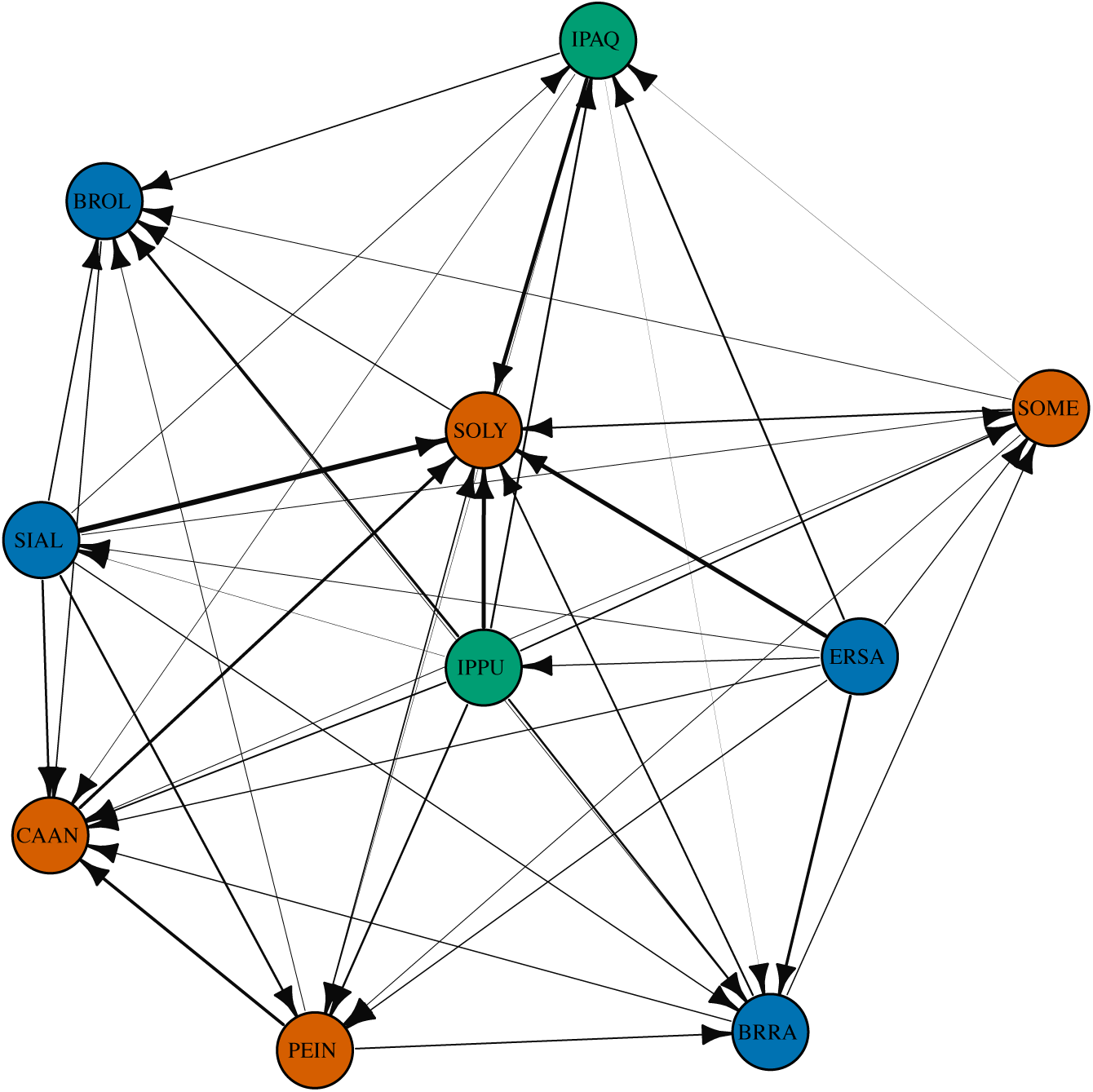
The lines connect the reciprocal pairwise comparison of effect sizes with 50% CP-50% HP. The edge of the arrow points to the species that loses, that is, the species with a larger effect size (more negative) in the reciprocal comparison. The species are coloured by family, blue (Brassicaceae), green (Convolvulaceae) and orange (Solanaceae). The intensity of effect is represented by the arrow’s size where greater difference between effect size correspond to a thicker line and vice versa. Species code: BROL: *Brassica oleracea*, BRRA: *Brassica rapa*, ERSA: *Eruca sativa*, SIAL: *Sinapis alba*, IPAQ: *Ipomoea aquatica*, IPPU: *Ipomoea purpurea*, CAAN: *Capsicum annuum*, PEIN: *Petunia integrifolia*, SOLY: *Solanum lycopersicum*, SOME: *Solanum melongena*.

### Phylogenetic distance

Effect sizes were not correlated with the phylogenetic distance matrix (Procrustes correlation = 0.57, sum of squares=0.68, *P* = 0.41). Analysis of phylogenetic signal of traits (**Table S7**) showed that only pollen size and stigmatic area had a strong significant phylogenetic signal (Pagel’s lambda = 0.99, *P* < 0.01; Pagel’s lambda = 0.86, *P* < 0.5 respectively).

### Plant functional traits

The species x species matrix of effect sizes was not correlated with the distance matrix of all traits (Euclidean distance) (Procrustes correlation = 0.58; sum of squares = 0.65, *P* = 0.63). Moreover, we did not find significant correlation between the species x species matrix of effect sizes and the distances matrix of each trait separately (**Table S8**).

Although the species x species matrix of effect sizes was not correlated with the traits when we analyze each family independently (Solanaceae and Brassicaceae) (Procrustes correlation = 0.79; sum of squares = 0.38, *P* = 0.75; Procrustes correlation = 0.60; sum of squares = 0.65, *P* = 0.85 respectively), significant correlations were identified when the analysis was conducted trait by trait (**Table S8**). For Solanaceae, the distance matrix of pollen-ovule ratio was significantly correlated with the species x species matrix of effect sizes (Procrustes correlation = 0.87; sum of squares = 0.25, *P* < 0.05) and for Brassicaceae, the distance matrix of style length was correlated with the species x species matrix of effect sizes (Procrustes correlation = 0.94; sum of squares = 0.12, *P* < 0.05).

The analysis of the absolute effect of traits on heterospecific pollen effect revealed a low/moderate correlation between traits and effect sizes (R^2^ = 0.25). Stigma size and pollen-ovule ratio was positively correlated with effect sizes (Estimates 0.53 +/- 0.11 and 0.30 +/- 0.11 respectively, *P* < 0.01). Moreover, donor pollen size interacted significantly with recipient stigma size (Estimate 0.25 +/- 0.11, *P* < 0.05) whereby larger pollen impacted more species with smaller stigmas and small pollen did not differ in effect sizes within the different ranges of stigma size (**Fig. 5**). However, effect size was not related to donor pollen size. Although we did not include style length in the model, this was highly correlated with stigma size. Hence, species with larger stigmas, higher pollen-ovule ratios and longer styles were less impacted by heterospecific pollen. Moreover, the additional test of the importance of donor and recipient identity on effect sizes as fixed effects showed that effect sizes depended significantly on recipient identity (Sum of squares = 67.01, *F*-value = 11.45, *P* < 0.01) but not on donor identity (Sum of squares = 5.40, *F*-value = 0.92, *P* = 0.5).

**Fig. 5.**
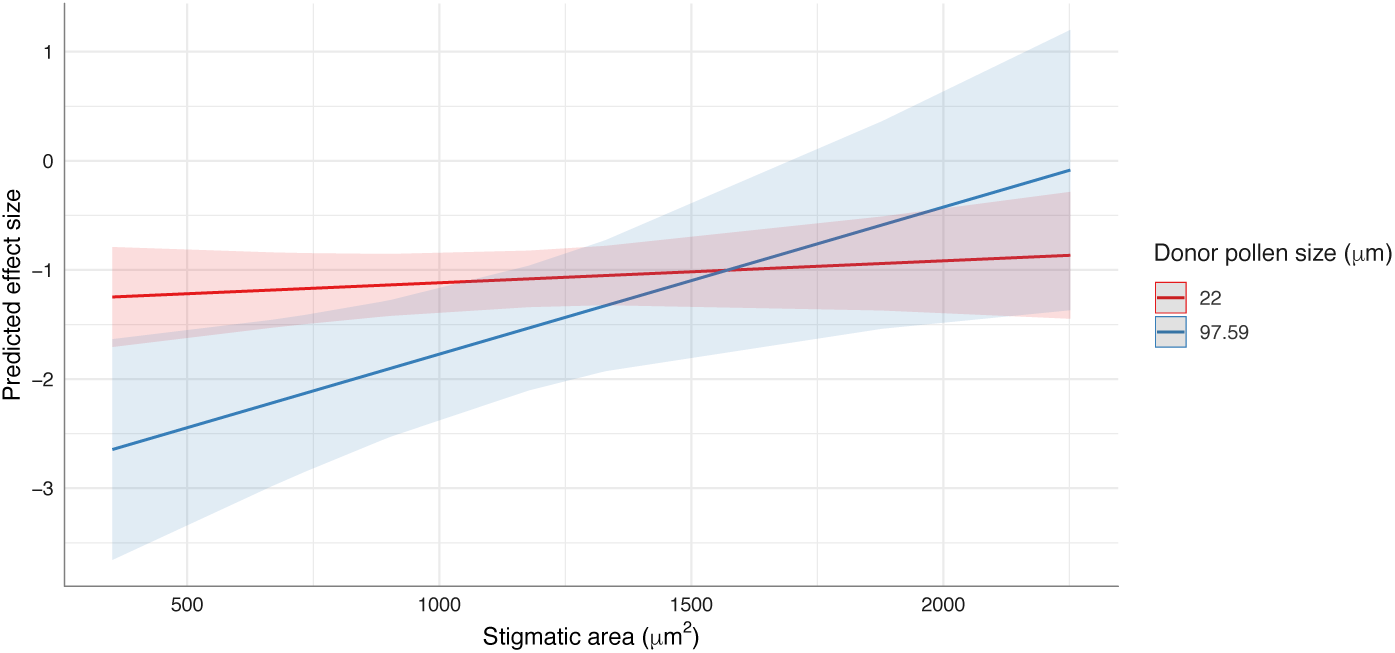
Plot of the interaction term of the stigmatic area and pollen size in the linear model of traits. The regression line coloured in red represents the correlation between the stigmatic area and the predicted effect size for small pollen, and the blue regression line for large pollen.

## DISCUSSION

Pollen transfer is an essential step for flowering plants that rely on sexual reproduction. However, pollinator inconstancy leads to the transfer of foreign pollen and therefore to the possibility of a negative impact on their fitness. As interactions can be asymmetrical and multifactorial, analyses of pairwise interactions are insufficient to encapsulate the broader range of multi-species interactions that occur, in particular how traits and phylogeny drive these interactions and effects (Arceo-Gómez *et al.*, 2016; Arceo-Gómez *et al.*, 2019). Here, we show the predominately negative impacts of heterospecific pollen are driven largely by recipient traits and the interaction between recipient-donor traits under specific circumstances of trait-matching. These traits and their asymmetries between species led to a hierarchical or transitive structure of pollen competition with clear winners and losers depending on specific combination of traits.

We found a clear hierarchical or transitive structure of pollen competition under approximately equal proportion of foreign pollen for all species, where 97.5% of the combinations of three species competed transitively. This study demonstrates that in an artificial co-flowering plant community, asymmetries in traits between species are an important determinant of plant reproductive success as a result of heterospecific pollen transfer. The average proportions of recipient/donor pollen found in Brassicaceae (80/20%), Solanaceae (60/40%) and Convolvulaceae (80/20%) led to a general detrimental effect for 67% of the treatments with very different outcomes between taxa, even with fixed quantities of pollen transferred in this experiment (Ashman & Arceo-Gómez, 2013; Arceo-Gómez *et al.*, 2019).

Style length, stigma size and pollen-ovule ratio were all correlated with heterospecific pollen effect size. First, we found that species with shorter styles were affected more by heterospecific pollen, as four of the five most affected species had shorter styles when compared with the rest of the species. These findings are consistent with previous pollination experiments conducted on closely related species which showed an asymmetrical growth of pollen tubes that depended on style length. Generally, the pollen of long styled species was able to grow the full length of short styled species but not vice versa (Lee, Page, McClure, & Holtsford, 2008; Tong & Huang, 2016). Second, we found that smaller stigmas are more likely to be impacted negatively by foreign pollen than larger stigmas (**Fig. 5**). Here, the species *S. lycopersicum,* which had the smallest stigma, was significantly more impacted by the larger pollen of Convolvulaceae family but not from the other two families with medium pollen size. This is likely because a smaller number of pollen grains can clog the stigmatic area easily and do not allow enough conspecific pollen grains to fertilize all the ovules. However, species with smaller stigmas could act as a filter in natural communities, reducing the heterospecific pollen load and therefore its impact (Montgomery & Rathcke, 2012). Further, while donor identity was of little importance to the outcome of the pollen-pistil interaction, the interaction between donor pollen size and stigmatic area had a significant impact on effect sizes (**Fig. 5**). Species with lower pollen-ovule ratios were also more likely to be impacted by foreign pollen. Pollen ovule-ratios are generally correlated with breeding system whereby low pollen-ovule ratios correspond to species with an autogamous breeding system and high pollen-ovule ratios to one that is xenogamous (Cruden, 1977). Therefore, species capable of selfing are more likely to suffer from fitness reduction by foreign pollen. This empirical evidence supports the predictive framework of Ashman & Arceo-Gómez (2013) for these specific traits, and highlights that studying recipient and donor traits separately may be insufficient (Arceo-Gómez *et al.*, 2019).

Given this study used fixed quantities of foreign pollen and detected a highly transitive network structure, it is possible that different quantities (and quality) of foreign pollen arrival in natural systems could result in varying competitive abilities among species. Yet recent studies have demonstrated that the competitive structure is likely to remain stable at community level over time unless the plant species involved develop strategies to cope with foreign pollen (Fang *et al.*, 2019). Therefore, in order to avoid exclusion through competitive dominance (Laird & Schamp, 2006), we hypothesize that the most vulnerable species are likely to develop strategies to minimize foreign pollen arrival, where species will adapt through tolerance or avoidance as pointed in previous studies (Ashman & Arceo-Gómez, 2013; Fang *et al.*, 2019).

The results of this study do not support the idea that self-incompatible species are less affected by heterospecific pollen (McClure *et al.*, 2011; Ashman & Arceo-Gómez 2013). In fact, we found contradictory results whereby the two *Brassica* species considered highly incompatible were highly susceptible to foreign pollen. We hypothesize that although foreign pollen recognition may take place for these species, it could lead to a stigmatic closure with important consequences for species fitness. As mixed mating systems are prevalent in nature (See **Fig.1** from Whitehead, Lanfear, Mitchell, & Karron, 2018), it is likely that foreign pollen arrival in addition to self-pollen is a common event. This could have important implications for species fitness because heterospecific pollen can have larger negative impacts with self pollination (Arceo-Gómez & Ashman, 2014). Further, few studies have investigated the impact of heterospecific pollen effect within wind pollinated species Martyniuk *et al.* 2015; Ikeru *et al.*, 2017).

In conclusion, trait asymmetries between species likely determine a transitive structure of competition with clear “winner” and “loser” species that share specific combinations of traits. We highlight that focusing on recipient traits, and particular trait matching interactions between donor and recipient species, could help to understand the mechanisms underlying the impacts of foreign pollen upon plant reproductive fitness. Here for the first time, we provide evidence of the competitive structure of heterospecific pollen with an artificial co-flowering community, with possibly important implications for plant coexistence and floral evolution.

## ACKNOWLEDGEMENTS

We gratefully acknowledge the help of G. Bible with the glasshouse work. We also thank J. Lobaton for his useful suggestions for lab work and help. Thanks are also due to O. Godoy for his useful advices for analysis. We also thanks S. Winkle for her help with plants and pollinations. Finally, JBL was supported by a University of New England postgraduate scholarship to carry out this work in Australia and RR was supported by an Australian Research Council Discovery Early Career Researcher Award DE170101349.

## AUTHORSHIP

JBL designed the study with input from all co-authors. JBL conducted the experiment and analyzed the data with IB contributions. JBL wrote the first draft of the manuscript and all authors contributed to the writing.

## DATA AVAILABILITY

All data and code are openly available at Zenodo https://doi.org/10.5281/zenodo.3601226.

