## Supporting Information for "Recipient and donor characteristics govern the hierarchical structure of heterospecific pollen competition networks"

Title: **Recipient and donor characteristics govern hierarchical structure in a heterospecific pollen competition network of co-flowering plants**

Authors: Jose B. Lanuza, Ignasi Bartomeus, Tia Lynn Ashman, Romina Rader

### Article acceptance date:

The following Supporting Information is available for this article:

**Table S1.** Species names, common names, varieties and sources of the different seeds.

**Table S2.** Numerical values of all the traits measured for each species.

**Table S3.** Seed set in percentage for hand cross-pollination, hand self-pollination, natural selfing and apomixis for all species.

**Table S4.** Species x species matrix with the significance of effect “yes” or “no” of the different donors on the seed set of the different recipient species.

**Table S5.** Estimates, standard error, *t*-value and *P*-value of the effect of the different 9 donors on each recipient species.

**Table S6.** Number of seeds produced with 100% foreign pollen treatments for the different recipient species.

**Table S7.** Phylogenetic signal and significance for all the different traits

**Table S8.** Procrustes analysis results.

**Figure S1.** Correlation matrix for all the different traits.

**Figure S2.** Total amount of pollen found for the different treatments.

**Figure S3.** Pollen ratios for the different recipient species.

**Figure S4** Pollen ratios for the different recipient species by family.

**Figure S5**. Statistical comparison of pollen ratios by family as pollen donor and recipient.

**Figure S6.** Violin plot of the reproductive biology of the species.

**Figure S7.** Grouped effect sizes by family for each recipient species.

| **Species** | **Common names** | **Variety** | **Source** |
| --- | --- | --- | --- |
| *Brassica oleracea* | Wild cabbage | Capitata | https://www.mrfothergills.com.au/ |
| *Brassica rapa* | Pak choi | Chinensis | https://www.mrfothergills.com.au/ |
| *Eruca sativa* | Rocket |  | https://www.mrfothergills.com.au/ |
| *Sinapis alba* | White mustard |  | https://www.mrfothergills.com.au/ |
| *Ipomoea aquatica* | Water spinach |  | https://www.theseedcollection.com.au/ |
| *Ipomoea purpurea* | Morning glory |  | http://www.shaman-australis.com.au |
| *Capsicum annuum* | Capsicum | California Wonder | https://www.edenseeds.com.au |
| *Petunia integrifolia* | Petunia |  | https://www.dianeseeds.com/ |
| *Solanum lycopersicum* | Tomato | Tommy Toe | https://www.mrfothergills.com.au/ |
| *Solanum melongena* | Eggplant | Little Fingers | https://www.4seasonsseeds.com.au/ |

**Table S1**. Species names, common names, varieties and sources of the different seeds

**Table S2.** Numerical values of all the traits measured for each species.

| **Spp/Traits** | **Pollen per anther** | **Pollen size (μm)** | **Number of ovules** | **Pollen-ovule ratio** | **Ovary length (μm)** | **Ovary width (μm)** | **Style length (μm)** | **Style width (μm)** | **Stigma length (μm)** | **Stigma width (μm)** | **Stigma area (μm^2^)** | **Selfing rate** | **SI index** |
| --- | --- | --- | --- | --- | --- | --- | --- | --- | --- | --- | --- | --- | --- |
| *Brassica oleracea* | 42033 | 27.72 | 29 | 8696.48 | 5927.46 | 1112.20 | 2324.53 | 650.95 | 527.81 | 884.88 | 622167.05 | 0.00 | 0.00 |
| *Brassica rapa* | 7133 | 25.35 | 26 | 1646.08 | 3531.94 | 877.76 | 1080.80 | 517.28 | 372.34 | 733.96 | 356582.77 | 0.00 | 0.00 |
| *Eruca sativa* | 22151 | 24.95 | 24 | 5537.75 | 4423.42 | 937.49 | 6595.94 | 730.84 | 733.24 | 668.01 | 345791.76 | 0.10 | 0.02 |
| *Sinapis alba* | 3507 | 33.59 | 6 | 3507.00 | 1981.32 | 1073.18 | 3624.95 | 773.86 | 631.04 | 906.31 | 548562.10 | 0.70 | 1.12 |
| *Ipomoea aquatica* | 858 | 70.10 | 4 | 1072.50 | 2384.95 | 1416.57 | 19441.30 | 453.06 | 1430.34 | 2251.70 | 3255360.52 | 0.60 | 0.75 |
| *Ipomoea purpurea* | 654 | 97.59 | 6 | 545.00 | 1056.54 | 1573.48 | 28230.08 | 579.81 | 1238.40 | 1878.23 | 2271922.22 | 1.00 | 2.74 |
| *Capsicum annuum* | 30761 | 32.46 | 241 | 765.83 | 3148.37 | 5797.99 | 3237.91 | 1057.22 | 724.83 | 1177.27 | 1060683.56 | 0.80 | 0.64 |
| *Petunia integrifolia* | 34657 | 24.74 | 220 | 787.66 | 3128.93 | 1771.59 | 14645.14 | 449.18 | 799.76 | 1322.44 | 1168415.58 | 0.90 | 0.26 |
| *Solanum lycopersicum* | 28915 | 22.00 | 92 | 1885.76 | 1161.33 | 1133.40 | 6466.81 | 310.19 | 190.22 | 351.60 | 91567.14 | 0.70 | 0.48 |
| *Solanum melongena* | 166989 | 25.18 | 1010 | 992.01 | 4020.49 | 3554.43 | 11329.30 | 936.53 | 961.83 | 1332.16 | 1137294.13 | 1.00 | 1.45 |

**Table S3.** Seed set in percentage for hand cross-pollination, hand self-pollination, natural selfing and apomixis for all species.

| **Species** | **Hand cross-pollination** | **Hand self-pollination** | **Natural selfing** | **Apomixis** |
| --- | --- | --- | --- | --- |
| *Brassica oleracea* | 32.07 | 0.00 | 0.00 | 0.00 |
| *Brassica rapa* | 44.97 | 0.00 | 0.00 | 0.00 |
| *Eruca sativa* | 23.75 | 0.41 | 0.00 | 0.00 |
| *Sinapis alba* | 43.33 | 48.33 | 5.00 | 15.00 |
| *Ipomoea aquatica* | 40.00 | 30.00 | 20.00 | 0.00 |
| *Ipomoea purpurea* | 31.67 | 86.66 | 31.67 | 0.00 |
| *Capsicum annuum* | 100.00 | 66.22 | 23.49 | 0.00 |
| *Petunia integrifolia* | 100.00 | 24.77 | 0.00 | 0.00 |
| *Solanum lycopersicum* | 90.38 | 43.48 | 70.00 | 0.00 |
| *Solanum melongena* | 60.48 | 87.97 | 21.56 | 0.00 |

|  | *B. oleracea* | *B. rapa* | *E. sativa* | *S. alba* | *I. aquatica* | *I. purpurea* | *C. annuum* | *P. integrifolia* | *S. lycopersicum* | *S. melongena* | Σ Significant effect of donors | % donors with significant effect |
| --- | --- | --- | --- | --- | --- | --- | --- | --- | --- | --- | --- | --- |
| *B. oleracea* |  | Yes | Yes | Yes | Yes | Yes | Yes | Yes | Yes | Yes | 9 | 100 |
| *B. rapa* | No |  | Yes | Yes | Yes | Yes | Yes | Yes | Yes | Yes | 8 | 88.9 |
| *E. sativa* | Yes | No |  | No | Yes | No | No | No | No | No | 1 | 22.2 |
| *S. alba* | No | Yes | No |  | Yes | Yes | Yes | No | No | No | 4 | 44.4 |
| *I. aquatica* | Yes | Yes | Yes | Yes |  | Yes | Yes | Yes | Yes | Yes | 9 | 100 |
| *I. purpurea* | No | No | Yes | No | No |  | Yes | No | Yes | No | 3 | 33.3 |
| *C. annuum* | Yes | Yes | No | Yes | No | Yes |  | Yes | Yes | Yes | 7 | 77.8 |
| *P. integrifolia* | Yes | No | No | Yes | Yes | Yes | No |  | Yes | Yes | 6 | 67.7 |
| *S. lycopersicum* | No | Yes | Yes | Yes | Yes | Yes | Yes | Yes |  | Yes | 8 | 88.9 |
| *S. melongena* | No | Yes | Yes | No | Yes | Yes | Yes | No | No |  | 5 | 55.6 |
|  |  |  |  |  |  |  |  |  |  |  | 60 | 66.7 |

**Table S4.** Species x species matrix with the significance of effect “yes” or “no” of the different donors on seed set from the linear mixed effect models. The category of “yes” represents significant effect of the donors (columns) on the different recipient species (rows) and “no” lack of significant reduction of seed set from the control. Significance was tested for all species by comparing with a control of hand cross-pollination with conspecific pollen.

**Table S5.** Output of linear mixed effect models of the effect of the different donors on the seed set of each recipient species.

|  | **Value** | **Std. Error** | **t-value** | **P-value** |
| --- | --- | --- | --- | --- |
| *B. oleracea* (intercept) | 1.993 | 0.222 | 8.982 | 0.000 |
| *B. oleracea ~ B. rapa* | -1.366 | 0.314 | -4.352 | 0.000 |
| *B. oleracea ~ C. annuum* | -0.866 | 0.314 | -2.760 | 0.007 |
| *B. oleracea ~ E. sativa* | -1.924 | 0.314 | -6.131 | 0.000 |
| *B. oleracea ~ I. aquatica* | -1.913 | 0.272 | -7.038 | 0.000 |
| *B. oleracea ~ I. purpurea* | -1.993 | 0.314 | -6.352 | 0.000 |
| *B. oleracea ~ P. integrifolia* | -1.635 | 0.314 | -5.210 | 0.000 |
| *B. oleracea ~ S. alba* | -1.566 | 0.314 | -4.989 | 0.000 |
| *B. oleracea ~ S. lycopersicum* | -1.474 | 0.314 | -4.697 | 0.000 |
| *B. oleracea ~ S. melongena* | -1.606 | 0.314 | -5.118 | 0.000 |
| *B. rapa* (intercept) | 2.370 | 0.190 | 12.492 | 0.000 |
| *B. rapa ~ B. oleracea* | -0.544 | 0.288 | -1.892 | 0.062 |
| *B. rapa ~ C. annuum* | -1.421 | 0.288 | -4.939 | 0.000 |
| *B. rapa ~ E. sativa* | -2.370 | 0.288 | -8.237 | 0.000 |
| *B. rapa ~ I. aquatica* | -1.918 | 0.288 | -6.669 | 0.000 |
| *B. rapa ~ I. purpurea* | -2.260 | 0.288 | -7.855 | 0.000 |
| *B. rapa ~ P. integrifolia* | -2.370 | 0.288 | -8.237 | 0.000 |
| *B. rapa ~ S. alba* | -2.370 | 0.288 | -8.237 | 0.000 |
| *B. rapa ~ S. lycopersicum* | -1.884 | 0.288 | -6.550 | 0.000 |
| *B. rapa ~ S. melongena* | -2.370 | 0.288 | -8.237 | 0.000 |
| *E. sativa* (intercept) | 1.336 | 0.352 | 3.802 | 0.000 |
| *E. sativa ~ B. oleracea* | -1.336 | 0.497 | -2.689 | 0.009 |
| *E. sativa ~ B. rapa* | 0.657 | 0.497 | 1.322 | 0.189 |
| *E. sativa ~ C. annuum* | -0.273 | 0.497 | -0.549 | 0.584 |
| *E. sativa ~ I. aquatica* | 0.870 | 0.497 | 1.749 | 0.084 |
| *E. sativa ~ I. purpurea* | -0.195 | 0.497 | -0.392 | 0.696 |
| *E. sativa ~ P. integrifolia* | 0.205 | 0.497 | 0.412 | 0.681 |
| *E. sativa ~ S. alba* | 0.324 | 0.497 | 0.652 | 0.516 |
| *E. sativa ~ S. lycopersicum* | -0.519 | 0.497 | -1.043 | 0.300 |
| *E. sativa ~ S. melongena* | 0.541 | 0.497 | 1.088 | 0.279 |
| *S. alba* (intercept) | 0.897 | 0.221 | 4.066 | 0.000 |
| *S. alba ~ B. oleracea* | -0.482 | 0.312 | -1.547 | 0.125 |
| *S. alba ~ B. rapa* | -0.897 | 0.312 | -2.875 | 0.005 |
| *S. alba ~ C. annuum* | 0.670 | 0.312 | 2.147 | 0.034 |
| *S. alba ~ E. sativa* | -0.541 | 0.312 | -1.735 | 0.086 |
| *S. alba ~ I. aquatica* | -0.626 | 0.312 | -2.007 | 0.048 |
| *S. alba ~ I. purpurea* | -0.718 | 0.312 | -2.300 | 0.024 |
| *S. alba ~ P. integrifolia* | 0.479 | 0.312 | 1.536 | 0.128 |
| *S. alba ~ S. lycopersicum* | 0.521 | 0.312 | 1.671 | 0.098 |
| *S. alba ~ S. melongena* | 0.199 | 0.312 | 0.639 | 0.524 |
| *I. aquatica* (intercept) | 0.797 | 0.081 | 9.816 | 0.000 |
| *I. aquatica ~ B. oleracea* | -0.797 | 0.115 | -6.941 | 0.000 |
| *I. aquatica ~ B. rapa* | -0.797 | 0.115 | -6.941 | 0.000 |
| *I. aquatica ~ C. annuum* | -0.797 | 0.115 | -6.941 | 0.000 |
| *I. aquatica ~ E. sativa* | -0.797 | 0.115 | -6.941 | 0.000 |
| *I. aquatica ~ I. purpurea* | -0.797 | 0.115 | -6.941 | 0.000 |
| *I. aquatica ~ P. integrifolia* | -0.797 | 0.115 | -6.941 | 0.000 |
| *I. aquatica ~ S. alba* | -0.589 | 0.115 | -5.129 | 0.000 |
| *I. aquatica ~ S. lycopersicum* | -0.797 | 0.115 | -6.941 | 0.000 |
| *I. aquatica ~ S. melongena* | -0.687 | 0.115 | -5.983 | 0.000 |
| *I. purpurea* (intercept) | 0.817 | 0.202 | 4.050 | 0.000 |
| *I. purpurea ~ B. oleracea* | 0.373 | 0.285 | 1.308 | 0.194 |
| *I. purpurea ~ B. rapa* | -0.407 | 0.285 | -1.429 | 0.157 |
| *I. purpurea ~ C. annuum* | -0.817 | 0.285 | -2.864 | 0.005 |
| *I. purpurea ~ E. sativa* | -0.748 | 0.285 | -2.621 | 0.010 |
| *I. purpurea ~ I. aquatica* | 0.409 | 0.285 | 1.436 | 0.155 |
| *I. purpurea ~ P. integrifolia* | -0.374 | 0.285 | -1.311 | 0.193 |
| *I. purpurea ~ S. alba* | -0.326 | 0.285 | -1.144 | 0.256 |
| *I. purpurea ~ S. lycopersicum* | -0.817 | 0.285 | -2.864 | 0.005 |
| *I. purpurea ~ S. melongena* | -0.235 | 0.285 | -0.824 | 0.412 |
| *C. annuum* (intercept) | 4.590 | 0.523 | 8.782 | 0.000 |
| *C. annuum ~ B. oleracea* | -1.799 | 0.739 | -2.434 | 0.017 |
| *C. annuum ~ B. rapa* | -2.505 | 0.739 | -3.389 | 0.001 |
| *C. annuum ~ E. sativa* | -0.628 | 0.739 | -0.849 | 0.398 |
| *C. annuum ~ I. aquatica* | -1.363 | 0.739 | -1.844 | 0.068 |
| *C. annuum ~ I. purpurea* | -3.996 | 0.739 | -5.406 | 0.000 |
| *C. annuum ~ P. integrifolia* | -4.590 | 0.739 | -6.210 | 0.000 |
| *C. annuum ~ S. alba* | -3.634 | 0.739 | -4.916 | 0.000 |
| *C. annuum ~ S. lycopersicum* | -1.521 | 0.739 | -2.058 | 0.042 |
| *C. annuum ~ S. melongena* | -3.295 | 0.739 | -4.457 | 0.000 |
| *P. integrifolia* (intercept) | 4.626 | 0.494 | 9.361 | 0.000 |
| *P. integrifolia ~ B. oleracea* | -1.844 | 0.812 | -2.271 | 0.025 |
| *P. integrifolia ~ B. rapa* | -0.965 | 0.812 | -1.189 | 0.237 |
| *P. integrifolia ~ C. annuum* | 0.783 | 0.812 | 0.965 | 0.337 |
| *P. integrifolia ~ E. sativa* | -0.911 | 0.812 | -1.122 | 0.265 |
| *P. integrifolia ~ I. aquatica* | -3.181 | 0.812 | -3.918 | 0.000 |
| *P. integrifolia ~ I. purpurea* | -4.213 | 0.812 | -5.189 | 0.000 |
| *P. integrifolia ~ S. alba* | -2.993 | 0.812 | -3.686 | 0.000 |
| *P. integrifolia ~ S. lycopersicum* | -2.403 | 0.812 | -2.959 | 0.004 |
| *P. integrifolia ~ S. melongena* | -2.658 | 0.812 | -3.273 | 0.001 |
| *S. lycopersicum* (intercept) | 4.391 | 0.237 | 18.542 | 0.000 |
| *S. lycopersicum ~ B. oleracea* | -0.467 | 0.410 | -1.137 | 0.258 |
| *S. lycopersicum ~ B. rapa* | -2.737 | 0.410 | -6.672 | 0.000 |
| *S. lycopersicum ~ C. annuum* | -4.391 | 0.410 | -10.705 | 0.000 |
| *S. lycopersicum ~ E. sativa* | -2.974 | 0.410 | -7.251 | 0.000 |
| *S. lycopersicum ~ I. aquatica* | -4.127 | 0.410 | -10.062 | 0.000 |
| *S. lycopersicum ~ I. purpurea* | -3.802 | 0.410 | -9.270 | 0.000 |
| *S. lycopersicum ~ P. integrifolia* | -1.823 | 0.410 | -4.445 | 0.000 |
| *S. lycopersicum ~ S. alba* | -3.722 | 0.410 | -9.076 | 0.000 |
| *S. lycopersicum ~ S. melongena* | -1.842 | 0.410 | -4.490 | 0.000 |
| *S. melongena* (intercept) | 6.342 | 0.698 | 9.091 | 0.000 |
| *S. melongena ~ B. oleracea* | -0.666 | 0.987 | -0.675 | 0.501 |
| *S. melongena ~ B. rapa* | -4.333 | 0.987 | -4.392 | 0.000 |
| *S. melongena ~ C. annuum* | -2.329 | 0.987 | -2.360 | 0.020 |
| *S. melongena ~ E. sativa* | -3.120 | 0.987 | -3.162 | 0.002 |
| *S. melongena ~ I. aquatica* | -6.342 | 0.987 | -6.428 | 0.000 |
| *S. melongena ~ I. purpurea* | -2.706 | 0.987 | -2.743 | 0.007 |
| *S. melongena ~ P. integrifolia* | -1.429 | 0.987 | -1.448 | 0.151 |
| *S. melongena ~ S. alba* | -1.107 | 0.987 | -1.122 | 0.265 |
| *S. melongena ~ S. lycopersicum* | -0.845 | 0.987 | -0.856 | 0.394 |

**Table S6.** Number of seeds produced with 100% foreign pollen treatments for the different recipient species. Recipient species with 0 seed production are not presented in the table.

| **Family** | **Recipient** | **Donor (100% pollen)** | **Seeds** |
| --- | --- | --- | --- |
| Brassicaceae | *B. oleracea* | *C. annuum* | 5 |
| Brassicaceae | *B. rapa* | *B. oleracea* | 2 |
| Brassicaceae | *B. rapa* | *B. oleracea* | 13 |
| Brassicaceae | *B. rapa* | *S. lycopersicum* | 1 |
| Brassicaceae | *B. rapa* | *B. oleracea* | 7 |
| Brassicaceae | *B. rapa* | *B. oleracea* | 5 |
| Brassicaceae | *S. alba* | *B. oleracea* | 7 |
| Brassicaceae | *E. sativa* | *C. annuum* | 6 |
| Brassicaceae | *E. sativa* | *C. annuum* | 1 |
| Solanaceae | *S. lycopersicum* | *S. alba* | 3 |
| Solanaceae | *S. melongena* | *P. integrifolia* | 36 |
| Solanaceae | *C. annuum* | *S. alba* | 127 |
| Solanaceae | *C. annuum* | *E. sativa* | 3 |

**Table S7.** Phylogenetic signal and significance for all the different traits.

| **Pagel’s lambda (λ)** | **Significance** | **Traits** |
| --- | --- | --- |
| 0.95 | 0.20 | Selfing rate |
| 0.99 | <0.01 | Pollen size |
| 6.61x10^-5^ | 1.00 | Pollen per anther |
| 6.61x10^-5^ | 1.00 | Number of ovules |
| 6.61x10^-5^ | 1.00 | Pollen-ovule ratio |
| 0.86 | <0.05 | Stigmatic area |
| 6.61x10^-5^ | 1.00 | Stigma length |
| 6.61x10^-5^ | 1.00 | Stigma width |
| 6.61x10^-5^ | 1.00 | Style length |
| 6.61x10^-5^ | 1.00 | Style width |
| 0.49 | 0.28 | Ovary length |
| 6.61x10^-5^ | 1.00 | Ovary width |
| 0.49 | 0.36 | SI index |

**Table S8.** Procrustes correlation, sum of squares and significance from Procrustes analysis between the matrix of effect sizes (species x species matrix) and the distance matrix of each trait for all families, just Solanaceae and just Brassicaceae.

| **Correlation** | **Sum of squares** | **Significance** | **Traits** | **Families** |
| --- | --- | --- | --- | --- |
| 0.34 | 0.88 | 0.74 | Selfing rate | All |
| 0.36 | 0.87 | 0.44 | Pollen size | All |
| 0.35 | 0.88 | 0.58 | Pollen per anther | All |
| 0.33 | 0.89 | 0.55 | Number of ovules | All |
| 0.30 | 0.91 | 0.86 | Pollen-ovule ratio | All |
| 0.37 | 0.86 | 0.67 | Stigmatic area | All |
| 0.62 | 0.62 | 0.10 | Stigma length | All |
| 0.46 | 0.79 | 0.46 | Stigma width | All |
| 0.43 | 0.82 | 0.43 | Style length | All |
| 0.59 | 0.65 | 0.12 | Style width | All |
| 0.34 | 0.88 | 0.58 | Ovary length | All |
| 0.35 | 0.88 | 0.94 | Ovary width | All |
| 0.49 | 0.76 | 0.23 | SI index | All |
| 0.86 | 0.26 | 0.25 | Selfing rate | Solanaceae |
| 0.76 | 0.42 | 0.33 | Pollen size | Solanaceae |
| 0.42 | 0.82 | 0.79 | Pollen per anther | Solanaceae |
| 0.52 | 0.73 | 0.83 | Number of ovules | Solanaceae |
| 0.87 | 0.25 | 0.04 | Pollen-ovule ratio | Solanaceae |
| 0.82 | 0.33 | 0.17 | Stigmatic area | Solanaceae |
| 0.89 | 0.20 | 0.08 | Stigma length | Solanaceae |
| 0.84 | 0.29 | 0.12 | Stigma width | Solanaceae |
| 0.78 | 0.39 | 0.50 | Style length | Solanaceae |
| 0.52 | 0.73 | 0.71 | Style width | Solanaceae |
| 0.64 | 0.60 | 0.79 | Ovary length | Solanaceae |
| 0.87 | 0.23 | 0.17 | Ovary width | Solanaceae |
| 0.50 | 0.75 | 1.00 | SI index | Solanaceae |
| 0.61 | 0.63 | 0.50 | Selfing rate | Solanaceae |
| 0.60 | 0.64 | 0.71 | Pollen size | Brassicaceae |
| 0.40 | 0.84 | 0.96 | Pollen per anther | Brassicaceae |
| 0.62 | 0.61 | 0.42 | Number of ovules | Brassicaceae |
| 0.49 | 0.76 | 0.96 | Pollen-ovule ratio | Brassicaceae |
| 0.55 | 0.70 | 0.58 | Stigmatic area | Brassicaceae |
| 0.92 | 0.15 | 0.08 | Stigma length | Brassicaceae |
| 0.63 | 0.60 | 0.38 | Stigma width | Brassicaceae |
| 0.94 | 0.12 | 0.04 | Style length | Brassicaceae |
| 0.79 | 0.37 | 0.33 | Style width | Brassicaceae |
| 0.59 | 0.65 | 0.50 | Ovary length | Brassicaceae |
| 0.54 | 0.70 | 1.00 | Ovary width | Brassicaceae |
| 0.50 | 0.75 | 0.83 | SI index | Brassicaceae |

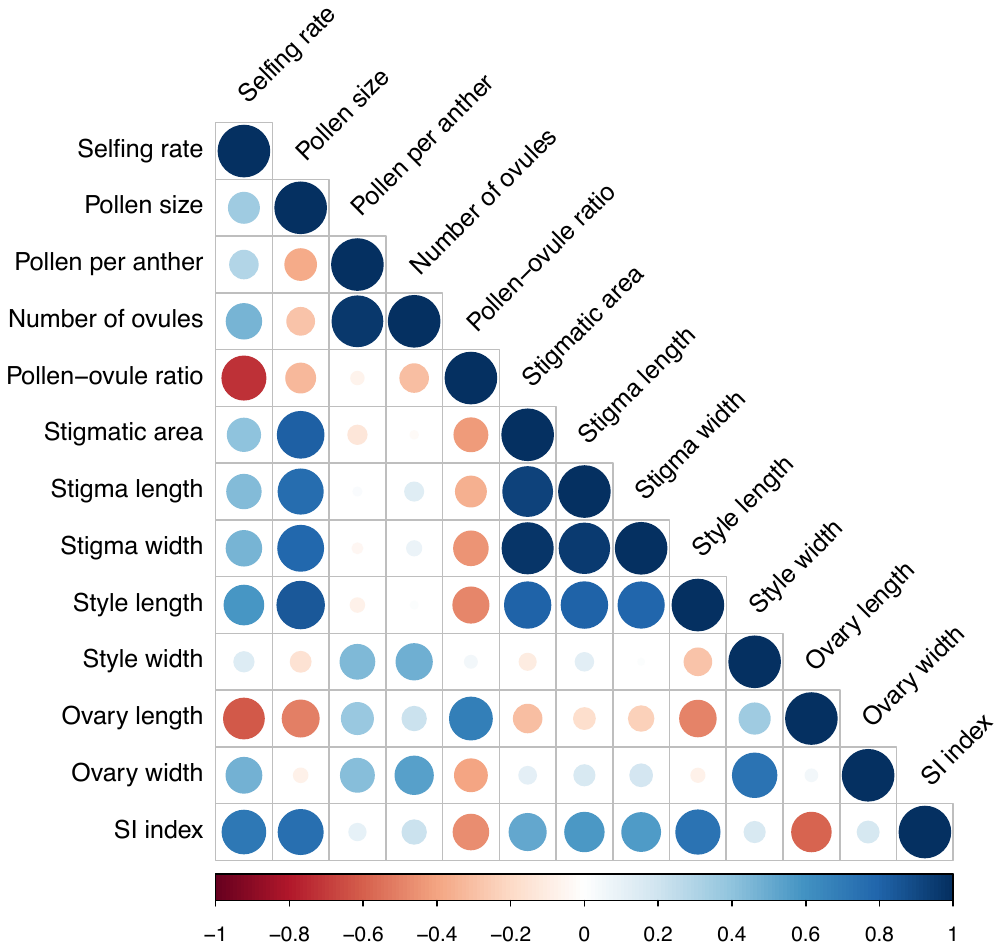

**Fig. S1** Graphical representation of the correlation matrix of the different reproductive traits considered in the experiment. Positive correlations are displayed in blue and negative in red with intensity and size of the circle proportional to the correlation coefficient from Pearson’s r.

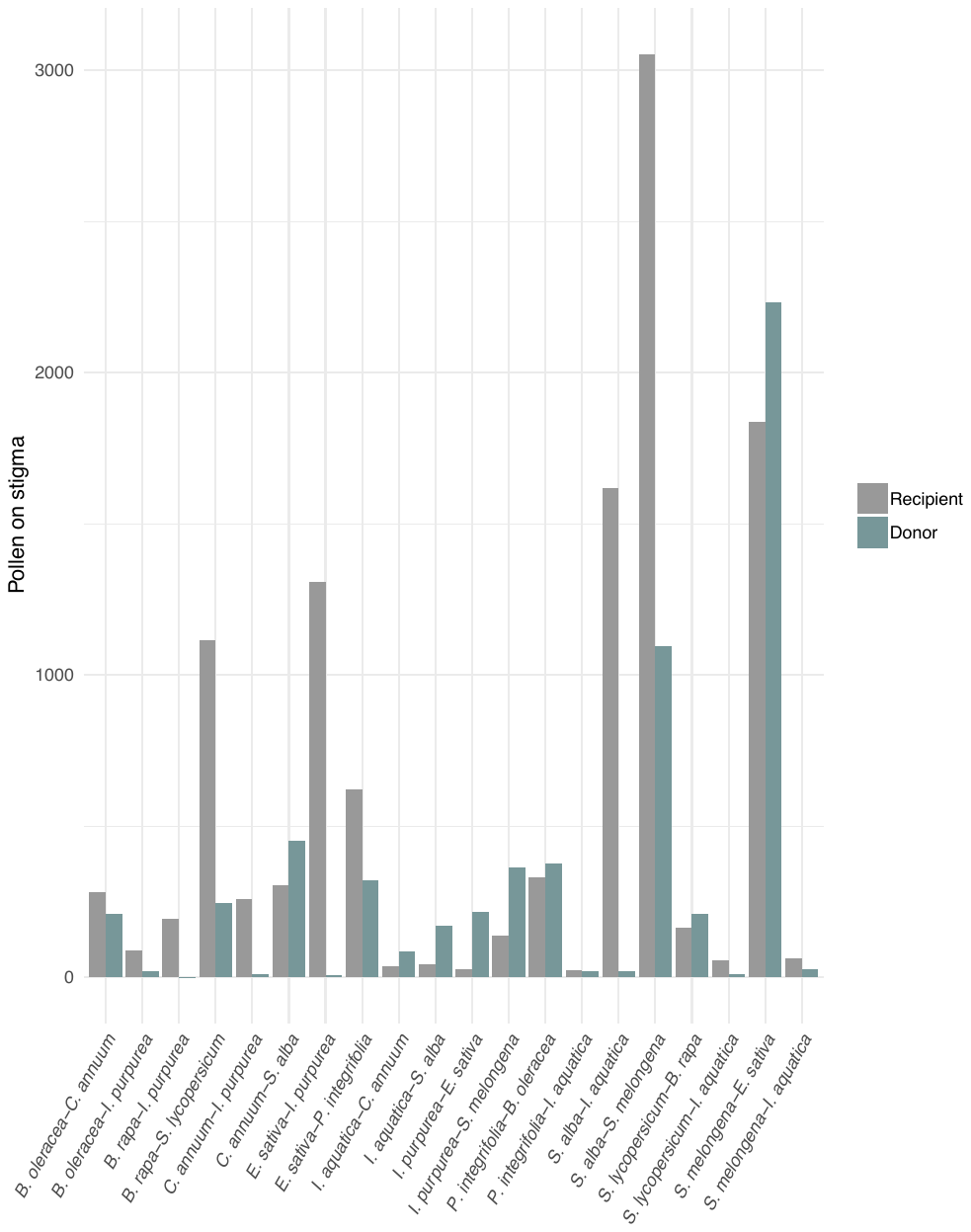

**Fig. S2** Total amount of pollen per stigma for 20 different treatments (see methods section for detailed explanation of the measurements). Each pair of bars correspond to the amount of focal pollen and donor pollen on the recipient species. The first species on the x-axis is the recipient species.

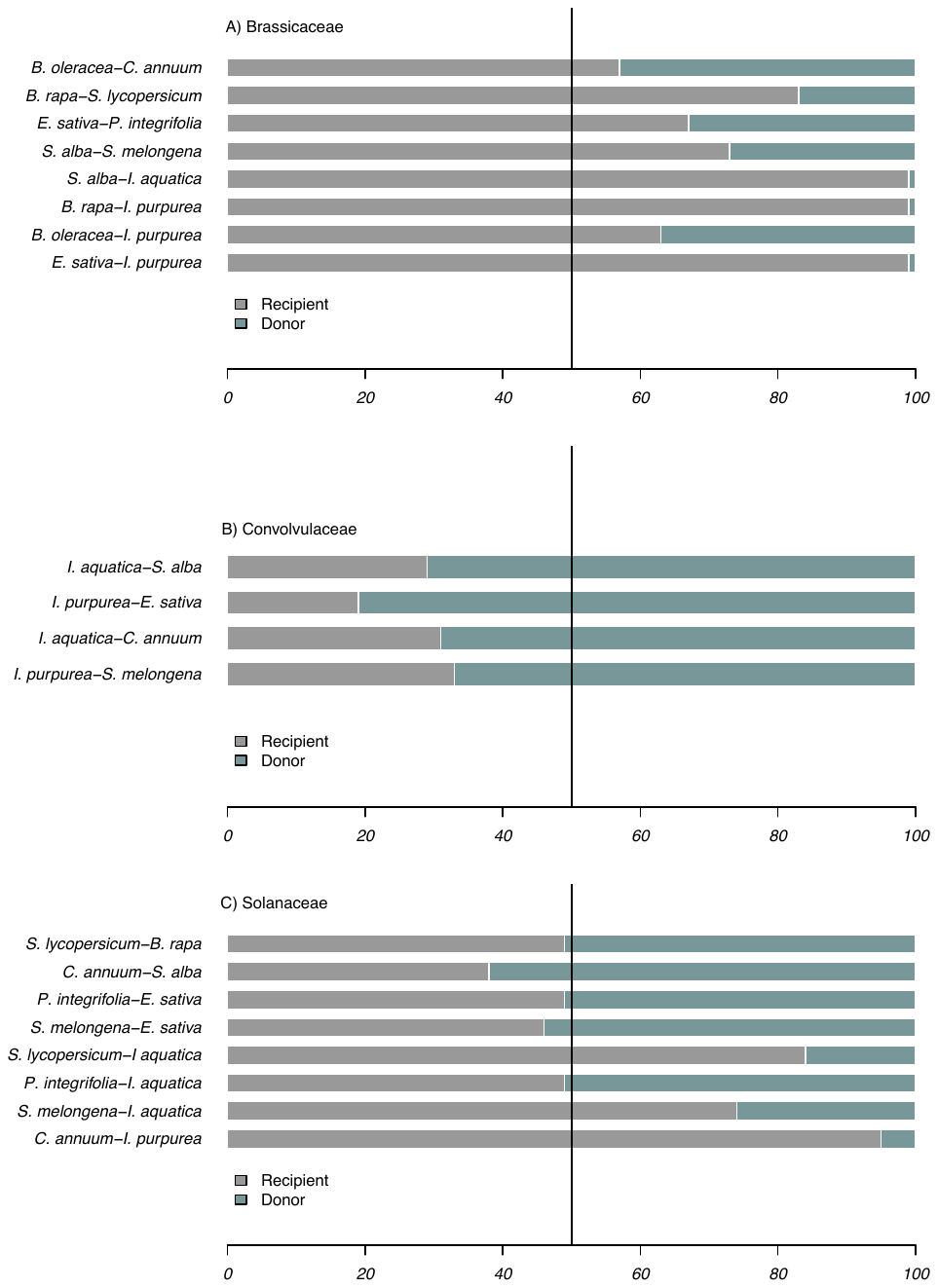

**Fig. S3** Pollen ratios from pollen counted on the stigma separated by family, A) Brassicaceae, B) Solanaceae and C) Convolvulaceae. The recipient and donor species appear coloured in grey and blue respectively. The vertical black line represents 50% pollen of both donor and recipient. The first species on the y-axis label is the recipient and the second the donor.

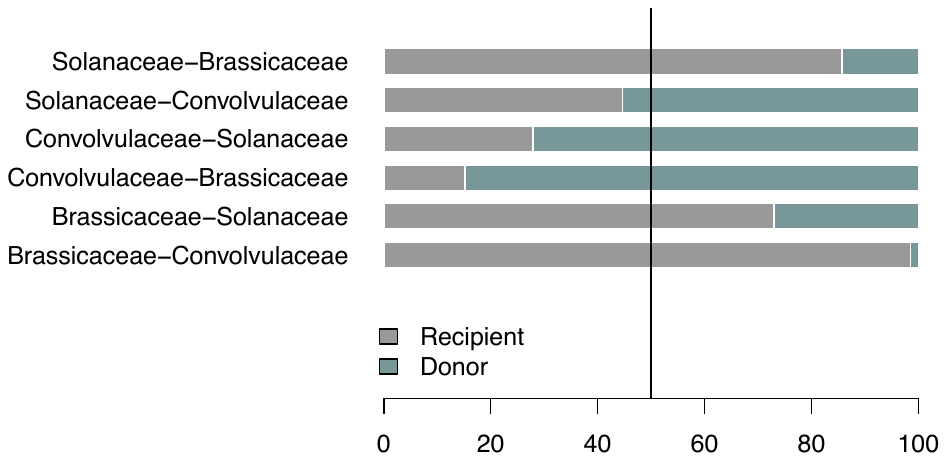

**Fig. S4** Average pollen ratios on stigma per family for 20 different treatments (see methods for treatment selection), where the first species on the y-axis is the recipient family and the second the donor. The vertical bar on intercept 50, represents equal proportions of both recipient (grey) and donor (light blue) pollen.

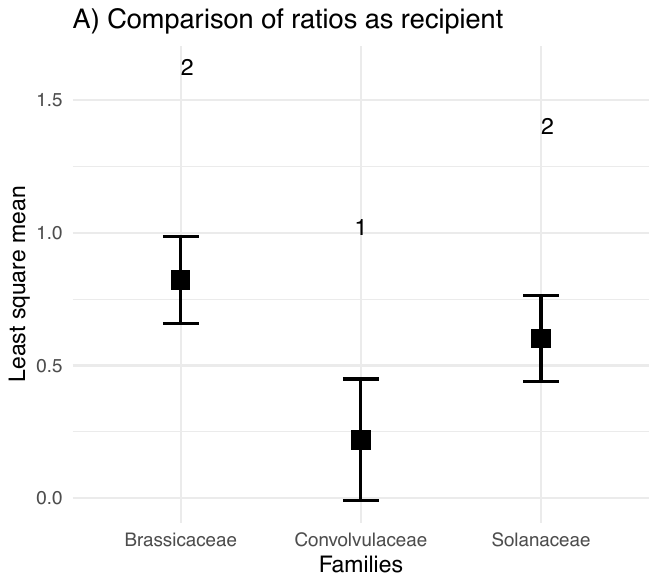

**Fig. S5** Pollen ratios comparisons between the different pollen recipient families where the boxes represent least square means, the error bars, confidence intervals 95%, and sharing numbers indicate no significant differences between groups (Tukey adjusted comparisons).

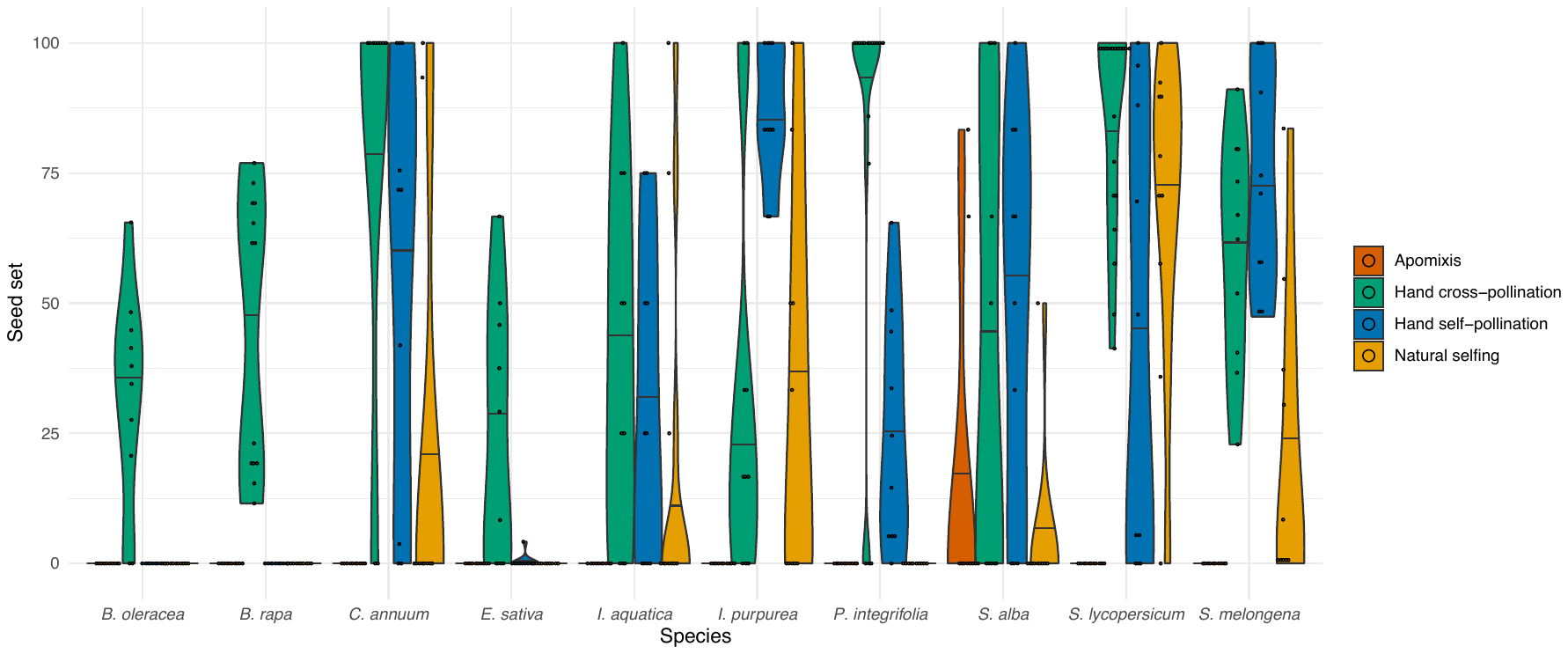

**Fig. S6** Violin plot showing the proportion seed set (%) of the ten species in response to each of four treatments (apomixis, hand cross pollination, hand self pollination and natural selfing). The coloured dots, represent the different values of seed set for each treatment.

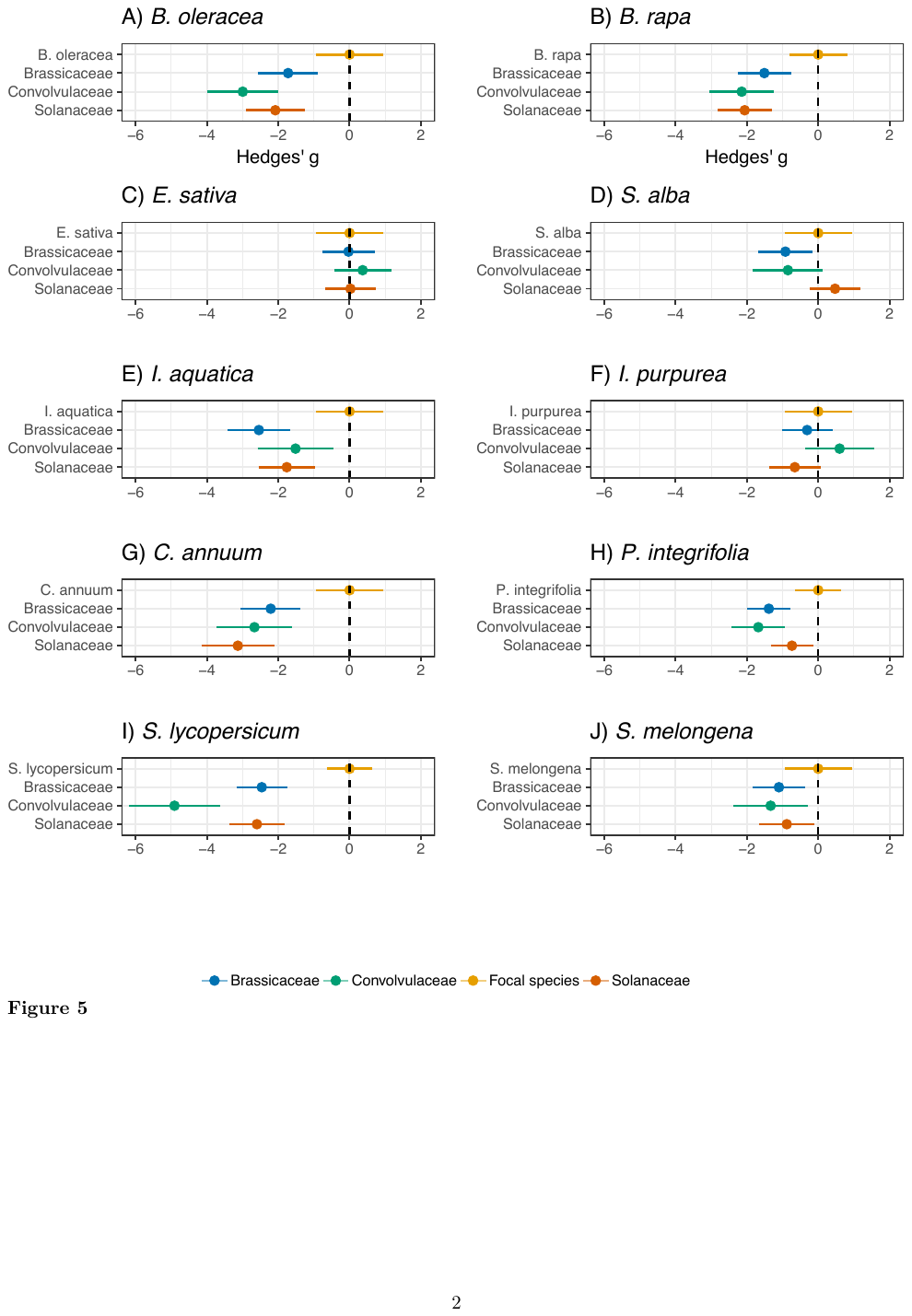

**Fig. S7** Effect sizes (95% confidence intervals) of the different families on each focal species (recipient) and a hand cross-pollination treatment (control). For each species the control treatment appears in yellow and the grouped effect per family in blue for Brassicaceae, green for Convolvulaceae and orange for Solanaceae.
